# ER calcium stores contribute to glucose-induced Ca^2+^ waves and intercellular connectivity in mouse pancreatic islets

**DOI:** 10.1101/2025.03.14.643121

**Authors:** Luis Fernando Delgadillo-Silva, Karen Dakessian, Guy A. Rutter

## Abstract

Defective insulin secretion is a hallmark of diabetes mellitus. Glucose-induced Ca^2+^ oscillations are critical for the stimulation of insulin secretion, though the mechanisms through which these propagate across the islet are poorly understood. Here, we use beta cell-targeted GCaMP6f to explore the role of endoplasmic reticulum (ER) Ca^2+^ mobilization in response to submaximal (11mM) and hyperglycemic (25mM) glucose concentrations. Inhibition of inositol 1,4,5 trisphosphate (IP_3_) receptors, and other ion channels, with 2-aminoethoxydiphenyl borate (2-APB) had minimal effects on the initial peak or intercellular connectivity provoked by 11mM glucose. However, 2-APB lowered subsequent glucose-induced cytosolic Ca^2+^ increases and connectivity at both 11 and 25mM glucose. Unexpectedly, the activation of IP_3_ receptors with the muscarinic acetylcholine receptor agonist carbachol had minimal impact on the initial peak elicited by 11 mM glucose, but Ca^2+^ waves at 11 and 25 mM glucose were more poorly coordinated. To determine whether ER calcium mobilization was sufficient to initiate Ca^2+^ waves we next blocked sarco(endo)plasmic Ca^2+^ ATPase (SERCA) pumps with thapsigargin, whilst preventing plasma membrane depolarization with the K_ATP_-channel opener, diazoxide. Under these conditions, an initial cytosolic Ca^2+^ increase was followed by secondary Ca^2+^ waves that slowly subsided. The application of carbachol alongside diazoxide still enhanced Ca^2+^ dynamics, though this activity was uncoordinated and beta cells were poorly connected. Our results show that ER Ca^2+^ mobilization plays a relatively minor role in the initiation and propagation of Ca^2+^ waves in response to glucose. On the other hand, ER stores are required to transition to sustained Ca^2+^ waves.

**Highlights:**

- IP3R inhibition or activation perturbs glucose-induced Ca^2+^ waves in islets
- ER store mobilization is insufficient to generate Ca^2+^ waves
- ER Ca^2+^ stores are required for sustained Ca^2+^ waves and beta cell connectivity

**GRAPHICAL ABSTRACT:** 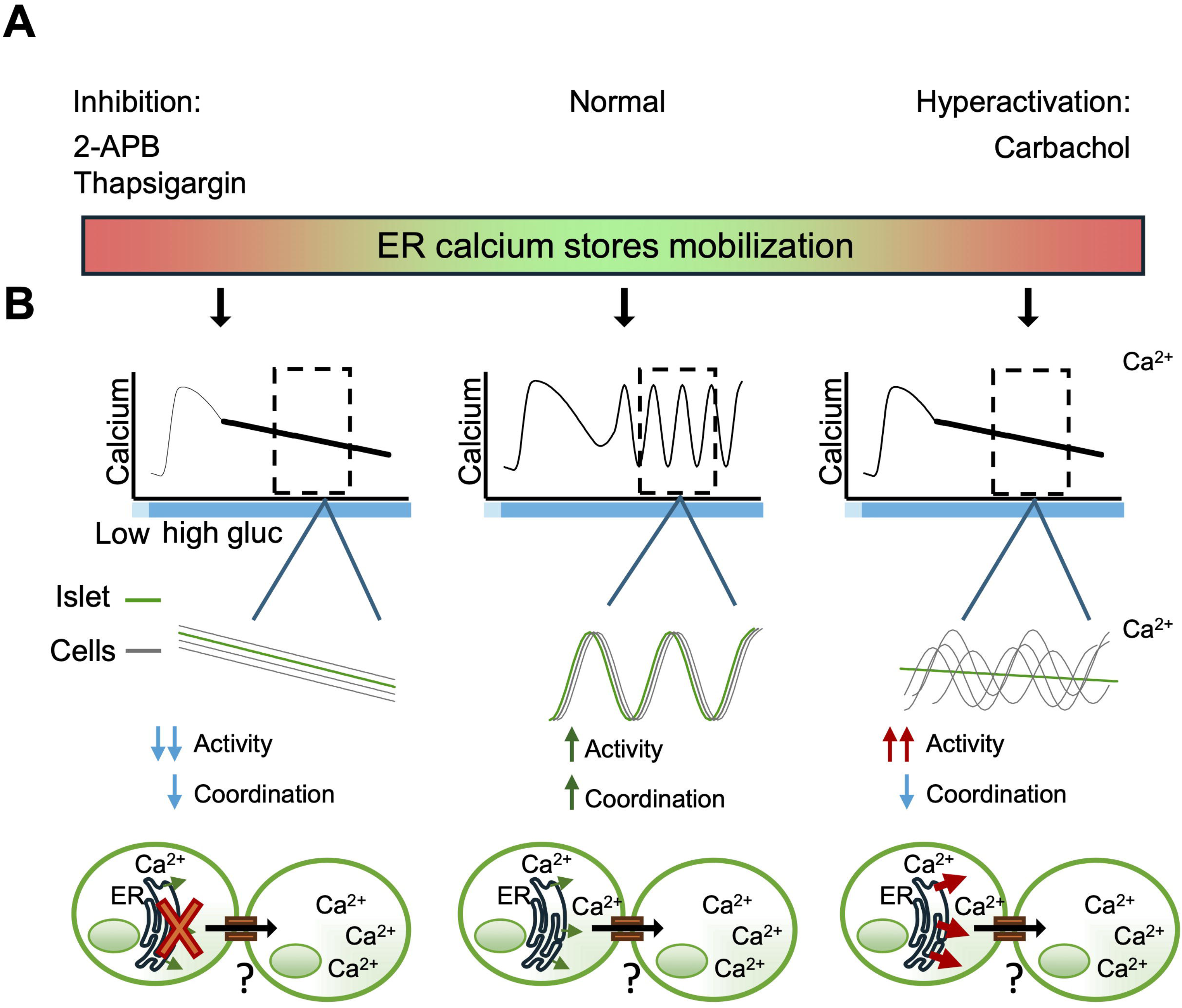

## 1. Introduction

Diabetes mellitus currently affects more than one in ten of the adult population (IDF). Whereas type 1 diabetes chiefly involves immune-mediated destruction of pancreatic islet beta cells [1], in the more prevalent type 2 diabetes beta cell dysfunction usually occurs in the face of insulin resistance [2]. Susceptibility to T2D involves both genetic factors, which chiefly impact beta cell function [3], and environmental challenges including obesity [4], which target insulin action and secretion.

In normoglycemic individuals, increases in circulating glucose enhance mitochondrial oxidative phosphorylation and increase the intracellular ATP/ADP ratio in the beta cell, leading to the closure of ATP-sensitive K^+^ (K_ATP_) channels, plasma membrane depolarisation and Ca^2+^ influx. Locally high concentrations of calcium [5] at the mouth of L-type voltage-gated calcium channels (VGCCs) then act on docked secretory granules [6] to cause exocytosis and the release of insulin [7]. Other coupling factors [8] and locally high ATP/ADP ratios [9] may also be involved (for discussion on this point see [10]), as well as multiple paracrine and neuronal inputs [11].

The role of intracellular Ca^2+^ mobilisation in the stimulation of insulin secretion by high glucose is usually thought to be relatively minor. Thus, blockade of external Ca^2+^ influx either through the use of diazoxide (to open K_ATP_ channels), or verapamil and other Ca^2+^ channel blockers [12] almost completely suppress insulin secretion in response to elevated glucose. Nevertheless, activation of G-protein (Gq)-coupled receptors including those for acetylcholine [13], and the opening of intracellular inositol 1,4,5-trisphosphate (IP_3_) receptors on the endoplasmic reticulum (ER) cause robust, if usually transient, cytosolic calcium increases, and potentiate glucose-stimulated insulin secretion [14]. Importantly, glucose-induced increases in intracellular IP_3_ are thought chiefly to be the result of Ca^2+^ activation of phospholipases, rather than serving as the initial driver of Ca^2+^ increases [15]. Nevertheless, recent evidence [16] has suggested a more active role for ER calcium stores in eliciting intracellular Ca^2+^ responses to glucose, and have suggested that ER-localized ryanodine receptors may also be involved.

In recent years, we [17–20] and several others [21–24] have shown that the pattern of intracellular Ca^2+^ changes across the islet is an important determinant of insulin secretion (reviewed in [25, 26]). Moreover, based on multicellular Ca^2+^ imaging across optical islet sections it has become evident that beta cells within the same islet display distinct roles. Thus, individual subgroups serve as “first responders”, “leaders” or highly-connected “hubs”. Inactivation of hubs [18], first responders [24] or leaders/first responders [19, 27] inhibits both the generation of Ca^2+^ oscillations and, in the case of hubs, insulin secretion.

The mechanisms involved in the generation of oscillations and waves in the islet, are nevertheless complex [26] and, at the level of individual beta cells, involve both metabolic and ionic components [28] [26]. The means by which the functional hierarchies above are established, and how oscillations are transmitted as spatially-organised waves, is also poorly understood. Potential roles for gap junctions [17, 29, 30], other cell types (e.g. delta cells [11, 20] and paracrine events [31], have all been proposed, whilst a recent report involving islets denuded of alpha and delta cells [32] has questioned the role of islet non-beta cells.

The notion that Ca^2+^ release from intracellular stores may be involved in generating oscillations and waves has not been explored in detail, though this process is critical in non-excitable cell types such as the liver, where IP3R-dependent calcium “flow” between cells underlies calcium waves [33]. In excitable cells, intracellular Ca^2+^ mobilisation may also lead to the opening of store-operated calcium channels [34] which in turn may modulate plasma membrane potential to fine tune the opening of VGCCs [35].

Given the above, it has seemed important to explore whether intracellular store opening may influence the generation and spread of spatially-organised Ca^2+^ oscillations across the islet using the highly speed multicellular imaging and analysis approaches described above. With this in mind, we have examined the effect on beta cell Ca^2+^ dynamics of manipulating ER Ca^2+^ release using multiple pharmacological approaches, including: (1) regulation of IP3R and other ion channel opening; (2) activation of Gq-coupled receptors and IP_3_ generation; (3) inhibition of Sarco(endo-) plasmic reticulum Ca^2+^ ATPases (SERCA). We show that each of these manoeuvres influences the dynamics of Ca^2+^ oscillation and wave generation, particularly under hyperglycemic conditions. Furthermore, our findings demonstrate roles for these stores in maintaining a functional hierarchy and connectivity between beta cells, which is likely to influence insulin secretion.

## 2. Materials and Methods

### 2.1. Animal Husbandry and Islet Isolation

All experimental manipulations were approved by the local ethical committee (CRCHUM, Montreal CIPA 2022–10,040 CM21022GRs). Colonies of Ins1Cre:GCaMP6^f/f^, in which the Ca^2+^ sensor GCaMP6f is expressed selectively in the beta cell [19,20], on a C57BL/6J background, were fed a regular chow diet and maintained at 21–23 °C, humidity 45–50 % and on a12 h day-night cycle.

Pancreatic islets were isolated from male mice aged between 8 and 16 weeks, as previously described [36, 37]. In brief, islets were collected in a Petri dish and hand-picked before being transferred to RPMI 1640 medium containing 11 mM glucose, 2 mM L-glutamine, 100 IU/mL penicillin, 100 μg/mL streptomycin, and fetal bovine serum (FBS - 10% v/v). Islets were cultured for 24 h at 37 °C in a humidified incubator with 5% CO_2_. For imaging, islets were transferred from the culture medium to a Krebs-HEPES-bicarbonate (KHB) buffer solution (130 mM NaCl, 3.6 mM KCl, 1.5 mM CaCl_2_, 0.5 mM MgSO_4_, 0.5 mM NaH_2_PO_4_, 24 mM NaHCO_3_, 10 mM HEPES; pH 7.4), initially at 3 mM glucose.

Imaging was performed essentially as per [20] using a Zeiss LSM 980 microscope with Airyscan 2, accompanied by a 37 °C incubation system. In brief, images were obtained using a 488 nm laser line for GCaMP6f excitation with a 40x objective for single cell resolution, and 20x or 2.5x for multiple islet imaging. Laser power was kept as low as possible (<1.5%) to minimize phototoxicity. All Ca^2+^ imaging videos were recorded using ZEISS Zen software across a single focal plane, with each plane recorded every 500 ms, providing a spatial resolution of 0.08 μm per pixel (512 x 512 pixels - 122.9 x 122.9 μm).

### 2.2. Drug Preparation

Diazoxide, 2-APB (2-Aminoethoxydiphenyl borate), and Carbachol were purchased from Sigma Aldrich, Tocris Bioscience, and Merck Emd Millipore, respectively. Stock solutions of Diazoxide, Thapsigargin, and 2-APB were prepared in DMSO and Carbachol in PBS. The concentrations used were as indicated in the figure legends. During live imaging, the drugs were added to the plate manually using a pipette.

### 2.3. Analysis of in vitro Beta Cell Imaging

Beta cell Ca^2+^ dynamics were quantified using the fluorescent signal derived from GCaMP6f. From each video, the islet and single cells were manually outlined using the ROI Manager in ImageJ (Fiji). The integrated fluorescence intensity of GCaMP6 was extracted then measured and exported into a CSV file. Integrated fluorescence intensity was normalized for the entire imaging duration using the following equation:

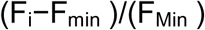

where Fi is the integrated fluorescence intensity at a given time. F_Min_ represents the minimum intensity recorded during live imaging. Cells that did not show an increase in GCaMP signal following KCl addition were not considered in the analysis. Single cell heat maps based on 2D analysis were created using GraphPad Prism 9.0 software, defining colors on a gradient from minimum to maximum value. The average GCaMP6f fluorescence unit per beta cell or per islet was normalized to the baseline, which is the minimum value recorded across the session. For better visualization, brightness and contrast were uniformly adjusted using the Brightness/Contrast tool in ImageJ.

### 2.4. Connectivity Analysis

Connectivity analyses were performed as previously described [20, 37]. In brief, smoothing was used to adjust Ca^2+^ signals via a moving average filter, contributing to 1% of the total length of the Ca^2+^ recording. Cell activity was represented in binary form, where any time point deviating 20% above the baseline is considered to be active, represented by “1”. Any inactive time point, under the threshold, is represented by a “0”. A minima-walking filter over 10% of the movie was applied to compensate from changes over time in the baseline. The following formula was used to calculate coactivity for each cell pair:

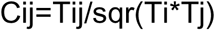

*Tij* denotes the total coactivity of each cell pair. *Ti* and *Tj* are the total activity time for the two cells compared, respectively.

Cells are considered to be linked if their t-test displayed a chance (>2 standard deviations i.e. p<0.01) probability within a corresponding distribution of the shuffled (10,000 times) binarized observed dataset, for the analyzed cell pairs.

A topographic representation of the connectivity was plotted in MATLAB (available from: https://www.mathworks.com/matlabcentral/fileexchange/24035-wgplot-weighted-graph-plot-a-better-version-of-gplot). Note that the edge colors indicate the strength of the coactivity between any two cells.

The analysis was divided into four frame sections: 3 mM (Baseline) from 0 to 360 frames (3 min), (11 mM) 361 to 2160 frames (15 min), (25 mM) 2161 to 4560 frames (20 min), and 40 mM KCl 4561 frames to 5040 (4 min). In each islet, cells with >70 % of connections having a connectivity of ≥0.8 were identified as highly-connected huns.

### 2.5 Statistics

Data are expressed as mean ± SD unless otherwise stated. Significance was tested by one or two-way ANOVA with Šidák or Brown-Forsythe multiple comparison tests, using GraphPad Prism 9 (GraphPad Software, San Diego, CA). *P* < 0.05 was considered significant.

## 3. Results

### 3.1. Inhibition of IP_3_R impairs glucose-induced intracellular Ca^2+^ dynamics and apparent beta cell-beta cell connectivity

To determine how intracellular Ca^2+^ mobilisation affects: (1) the generation of calcium oscillations and waves in the islet, (2) the appearance of functional sub-groups of cells, and (3) apparent cell-cell connectivity, we used islets from mice in which the genetically-encoded Ca^2+^ sensor GCaMP6f is expressed exclusively in the cytosol of the pancreatic beta cell [19, 37] (Fig 1A). We performed a glucose ramp from 3 to 11 and 25 mM glucose allowing 15 min. per condition (Fig 1B i) 11mM and ii) 25mM). Stepping from 3 to 11 mM glucose, that latter representing a stimulatory but sub-maximal concentration of the sugar, typically caused an initial increase in intracellular Ca^2+^, when assessed across the islet, followed by Ca^2+^ oscillations with a period of 1-2 min (movie 1). This frequency was further increased at 25 mM glucose (Fig 1C-D, Supp. Fig 1). The latter represents a near maximal concentration of the sugar with respect to insulin secretion, and is usually only observed in vivo in subjects with diabetes [38]. Finally, KCl was used to depolarise the plasma membrane and open VGCCs. Beta cell-beta cell connectivity (Fig 1D i) 11mM, ii) 25 mM) was assessed by co-activity measurements (Methods, *2.4*) and indicated a high degree of synchrony at each glucose concentration. Particularly highly connected (“hub”) cells, as identified after data binarization and assessment of co-activity [18–20] are indicated as black dots.

**Figure 1.**
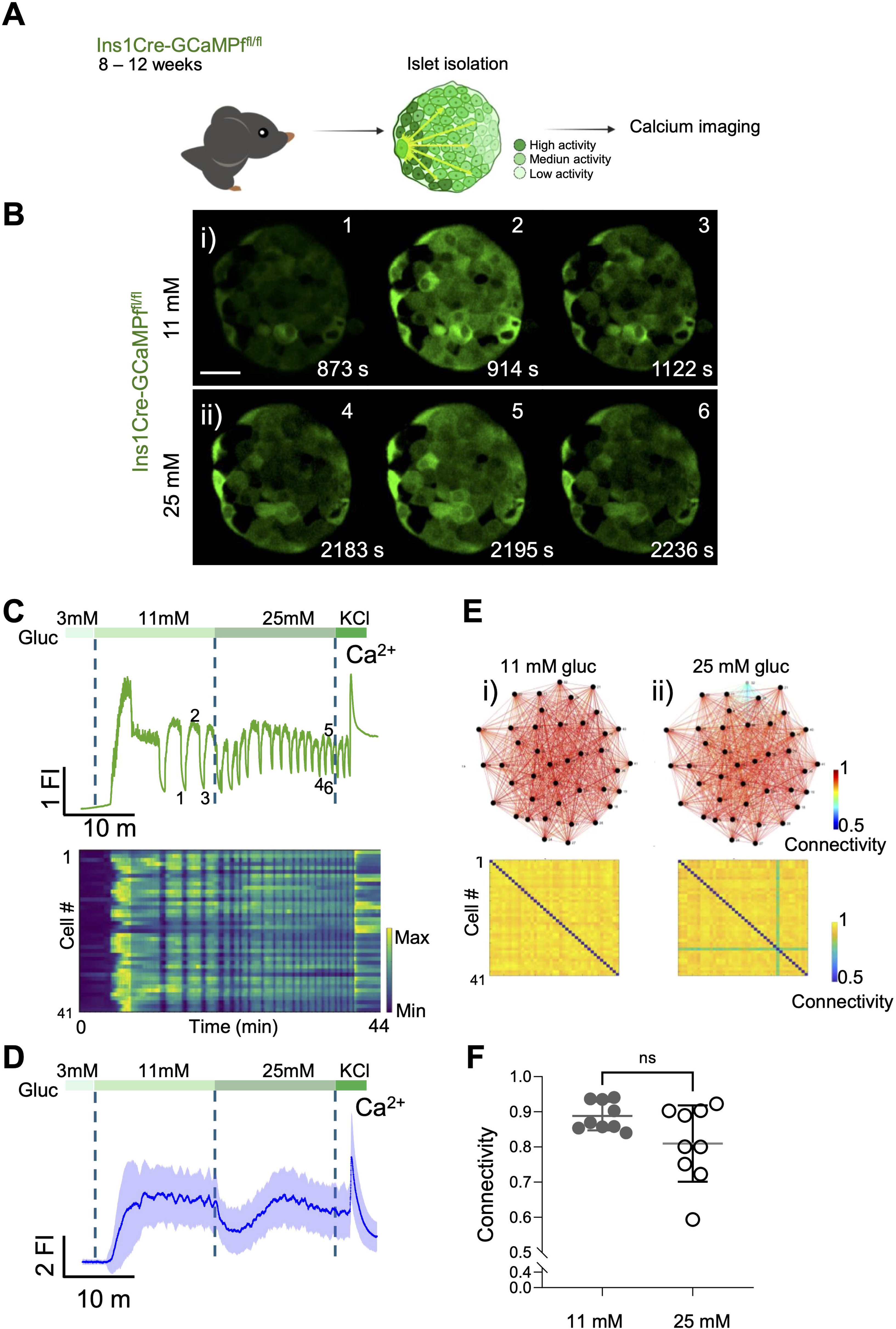
Glucose-stimulated Ca^2+^ Influx Imaged *in vitro* from mouse islet expressing GCaMP6f. **A)** Schematic representation of an isolated islet from a transgenic mouse expressing the genetically encoded Ca^2+^ indicator GCaMP6 (green), specifically in pancreatic beta-cells. GCaMP6 allows the study of glucose-induced Ca2+ entry into beta cells, detected through changes in green fluorescence dependent on intracellular Ca2+ concentration. **B)** Series of confocal calcium imaging of pancreatic islet showing GCaMP6f signal specifically in beta-cells at the indicated time points (in sec.) during calcium waves oscillations at **i)** 11 mM and **ii)** 25 mM glucose. Scale bar = 25 µm. **C)** Fluorescent trace of the GCaMP6 signal from the islet shown in **(B)** demonstrating calcium influx during the glucose ramp (3 mM, 11 mM, 25 mM) and membrane depolarization induced by 40 mM KCl. The raster plot corresponds to the signal from individual cells. **D)** Average calcium fluorescent traces from islets exposed to a glucose ramp (3 mM, 11 mM, 25 mM) and membrane depolarization by 40 mM KCl (n=3 WT 75-islets). The blue line shows the average GCaMP6f fluorescent and the light-blue the standard deviations from the imaged islets. **E)** Connectivity map and matrix of the islet shown in **(B)** at **i)** 11 mM and **ii)** 25 mM glucose. **F)** Connectivity from individual islets n=4 WT 9-islets (unpaired 2-tailed Student’s t-test. ns= not significant).

2-Aminoethoxydiphenyl borate (2-APB) is a low selectivity inhibitor of IP_3_ receptors (IP_3_R) which also exerts effects on sarco(endo)plasmic Ca^2+^ATPase (SERCA) pumps and ER leak activities [39–41]. As shown in Fig 2A and Supp movie 2, 2-APB had no impact on the initial response to an increase in glucose from 3 to 11 mM, but suppressed the generation of subsequent Ca^2+^ oscillations and the response to a further rise in glucose to 25 mM (Fig 2 B-C). Confirming the efficacy of the drug at this concentration, 2-APB largely abolished the response to the acetyl choline (AcCh) receptor analogue, carbachol, used as an agonist at metabotropic, Gq-coupled AcCh receptors (data not shown). Quantitation of these data (Supp. Fig 1) revealed tendencies towards a decrease in the amplitude of the oscillations at 11 mM glucose (Supp Fig 1B-C), with no effect on average peak width or frequency. On the other hand, all three parameters were significantly lowered by 2-APB during treatment at 25 mM glucose (Supp Fig 1D). Similar observations were main when 2-APB was added shortly after the initial step increase to 11 mM glucose (Supp. Fig. 3 and Movie 2).

**Figure 2.**
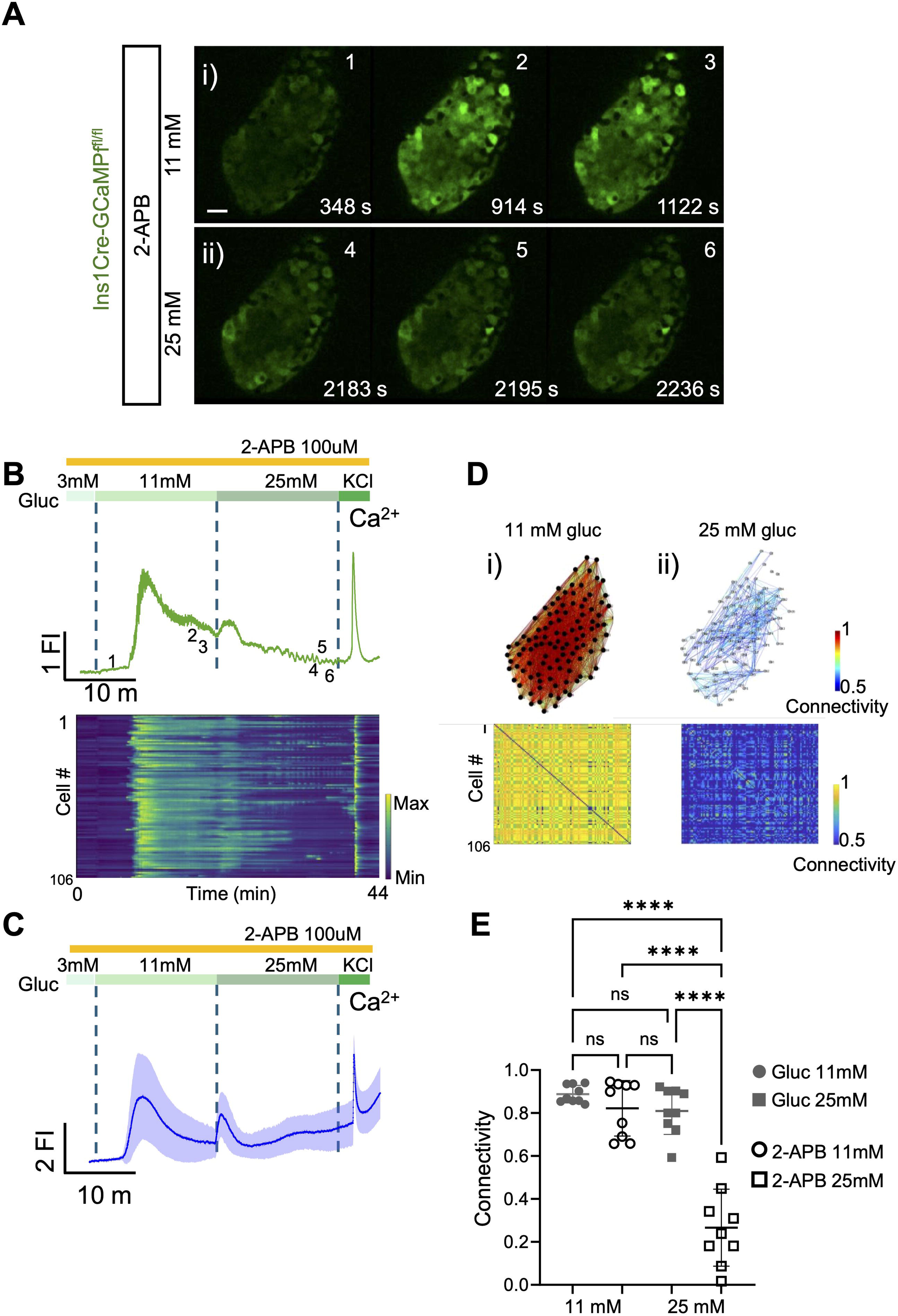
Pharmacological inhibition of IP_3_R with 2-APB impairs glucose-induced intracellular Ca^2+^ dynamics and cell-cell connectivity. **A)** Series of confocal calcium imaging of pancreatic islet at the indicated time points (in sec.), exposed to the IP3-inhibitor 2-Aminoethoxydiphenyl borate (2-APB) at **i)** 11 mM and **ii)** 25 mM glucose. Scale bar = 25 µm. **B)** Fluorescent trace of the GCaMP6 signal from the islet shown in **(A)** during the glucose ramp (3 mM, 11 mM, 25 mM) and membrane depolarization by 40 mM KCl. The raster plot corresponds to the signal from individual cells. **C)** Average calcium fluorescent traces from islets exposed to 2-APB during a glucose ramp (3 mM, 11 mM, 25 mM) and membrane depolarization by 40 mM KCl. The blue line shows the average GCaMP6f fluorescent and the light-blue the standard deviations from the imaged islets (n=4 WT 103-islets). **D)** Connectivity map and matrix of the islet shown in **(A)** at **i)** 11 mM and **ii)** 25 mM glucose. **E)** Connectivity from individual islets (glucose n=4 WT 9-islets; 2-APB n=4 WT 9-islets) (unpaired 2-tailed Student’s t-test. ns= not significant, **** *P*<0.0001).

Examined in individual islets (Fig 2D i)) beta cell-beta cell connectivity at 11 mM glucose in the presence of 2-APB was not significantly different from that in control conditions, although oscillations tended to be observed less often in the presence of the inhibitor. On the other hand, at 25 mM glucose, 2-APB caused a drastic lowering of connectivity (Fig. 2D ii)) compared to control (no drug) conditions (Fig. 2E) and consequently a sharp lowering in the number of highly connected “hub” cells (Fig. 2Dii versus Fig. 1Eii). Thus, at 11mM glucose, and average of 94.8% ± 8.5 were identified as “hubs”, similarly, at 25mM and average of 79.6% ± 32.2 were hubs. However, in 2-APB treated islets, an average of 60.1% ± 46.1 were identified as “hubs” at 11 mM glucose, whereas at 25mM no “hub” cells were detected under these conditions.

### 3.2. Muscarinic receptor activation generates exaggerated intracellular Ca^2+^ increases but does not impair cell-cell connectivity

In order to explore the impact of IP_3_-mediated ER calcium mobilisation on beta cell Ca^2+^ dynamics islet-wide, we used carbachol to activate metabotropic Gq-coupled AcCh receptors, leading to the activation of phospholipase C. Treatment with carbachol 60 s before a step increase in glucose (3 to 11 mM) cause a substantial increase in cytosolic Ca^2+^, followed by vigorous baseline oscillations (Fig 3A(i)-A(ii) and Movie 3). As expected, and indicating that glucose-induced calcium increased chiefly involve voltage-dependent influx across the plasma membrane, these oscillations were almost completely suppressed by the K_ATP_ channel opener, diazoxide (Fig 3A(iii)-A(vi) and Movie 4). Nevertheless, after treatment with CCh, 11 mM glucose elicited small but detectable increases across the islet, even in when diazoxide was present (Fig 3A(v)-A(vi) and Movie 5) consistent with actions of glucose which are independent of membrane depolarisation and VGCC-mediated Ca^2+^ influx.

**Figure 3.**
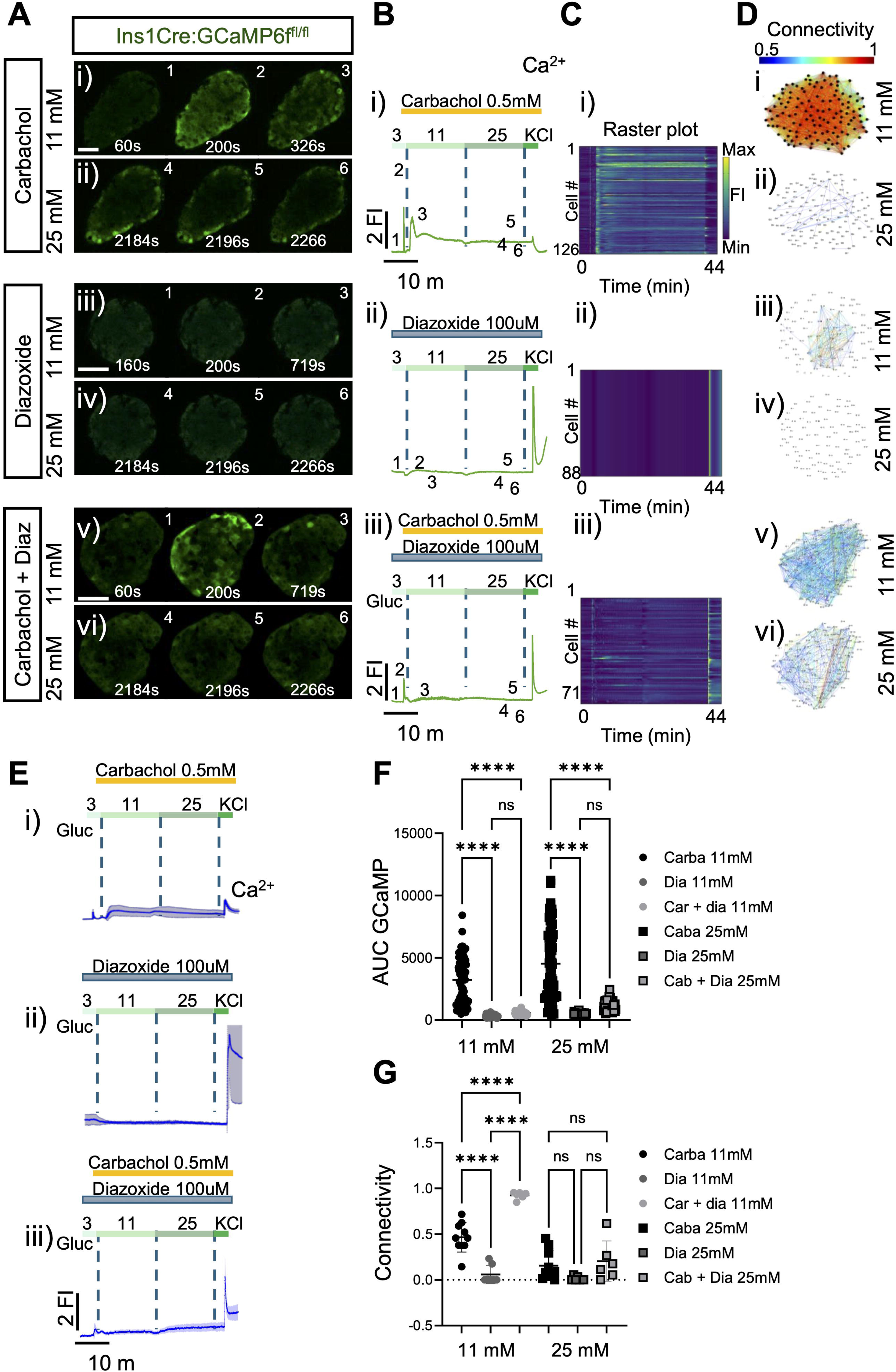
Muscarinic receptor agonism by carbachol induces calcium transients but impairs cell-cell connectivity. **A)** Series of confocal calcium imaging of pancreatic islet at the indicated time points (in sec.), exposed to the muscarine receptor agonist carbachol, to the K_ATP_ channel opener, diazoxide, which inhibits membrane depolarization and its combination (carbachol+diazoxide) at **i)** 11 mM and **ii)** 25 mM glucose. Scale bar = 25 µm. **B)** Fluorescent traces of the GCaMP6 signal from the islets shown in **(A)** during the glucose ramp (3 mM, 11 mM, 25 mM) and membrane depolarization by 40 mM KCl. **C)** Raster plot corresponds to the signal from individual cells from the islets shown in **(A)**. **D)** Connectivity maps of the islets shown in **(A)** at **i)** 11 mM and **ii)** 25 mM glucose. **E)** Average calcium fluorescent traces from islets exposed to carbachol, diazoxide or cabachol+diazoxide during a glucose ramp (3 mM, 11 mM, 25 mM) and membrane depolarization by 40 mM KCl. **F)** Calcium AUC quantifications from carbachol, diazoxide or carbachol and its combination at 11mM and 25mM glucose. Each symbol represents an islet (2-way paired ANOVA, Tukey’s correction \*\*\*\**P* ≤ .0001, ns, not significant; n=3 WT 71-islets for carbachol, n=3 WT 45-islets for diazoxide and n=3 WT 47-islets from diazoxide+carbachol). **G)** Connectivity quantifications from carbachol, diazoxide and carbachol+diazoxide at 11mM and 25mM glucose. Of note, at 25mM glucose, islets exposed to carbachol showed poorly coordinated activity, leading to impaired connectivity. Each symbol represents an islet (2-way paired ANOVA, Tukey’s correction \*\*\*\**P* ≤ .0001, ns, not significant; n=4 WT 10-islets for carbachol, n=3 WT 7-islets for diazoxide and n=3 WT 6-islets from diazoxide+carbachol).

At 11 mM glucose, carbachol caused only a minor decrease in connectivity *versus* that observed in the absence of the agonist (0.85 to 0.5; Fig 3D(i) versus Fig 1E(i)). On the other hand, overall Ca^2+^ dynamics, cell-cell connectivity and hub cell number at 11 mM glucose alone were low in the presence of diazoxide, as expected (Fig 3D(iii)). In the additional presence of CCh, connectivity in the presence of 11 mM glucose was detectably increased (Fig 3A(v), B(iii), C(iii), D(v)). In contrast, at 25 mM glucose, connectivity was low irrespective of the presence of diazoxide or CCh (Fig 3D(ii), D(iv), D(vi) versus Fig 1 E(i), E(ii)).

### 3.3 ER store depletion after SERCA pump inhibition leads to sustained Ca^2+^ increases, but barely impairs connectivity

The preceding experiments provided evidence that intracellular Ca^2+^ mobilisation may contribute to the establishment and propagation of Ca^2+^ waves, at last at high glucose concentrations. To further explore this possibility, we next explored the generation of islet Ca^2+^ waves after depleting Ca^2+^ stores with the SERCA pump inhibitor, thapsigargin. We depleted the ER calcium stores by preincubating the islets for 1h in 10uM thapsigargin. In the presence of the inhibitor, 11 m glucose-induced increases in cytosolic Ca^2+^ were exaggerated compared to those in control conditions (Fig 4A(i), B(i), E versus Fig 1B-D), but baseline oscillations and waves were rarely observed (Fig. 4A and Movie 6). As shown earlier (Fig. 3A(iii), A(iv), E(ii), glucose (11 or 25 mM) - induced Ca^2+^ oscillations were completely blocked in the presence of diazoxide, yet thapsigargin addition still elicited a Ca^2+^ peak (reflecting the acute inhibition of SERCA pumps). Under these conditions, a subsequent increase to 25 mM glucose caused clear, if relatively brief and poorly coordinated, Ca^2+^ oscillations (Fig 4A(iii), A(iv), E(ii) and Movie 7).

**Figure 4.**
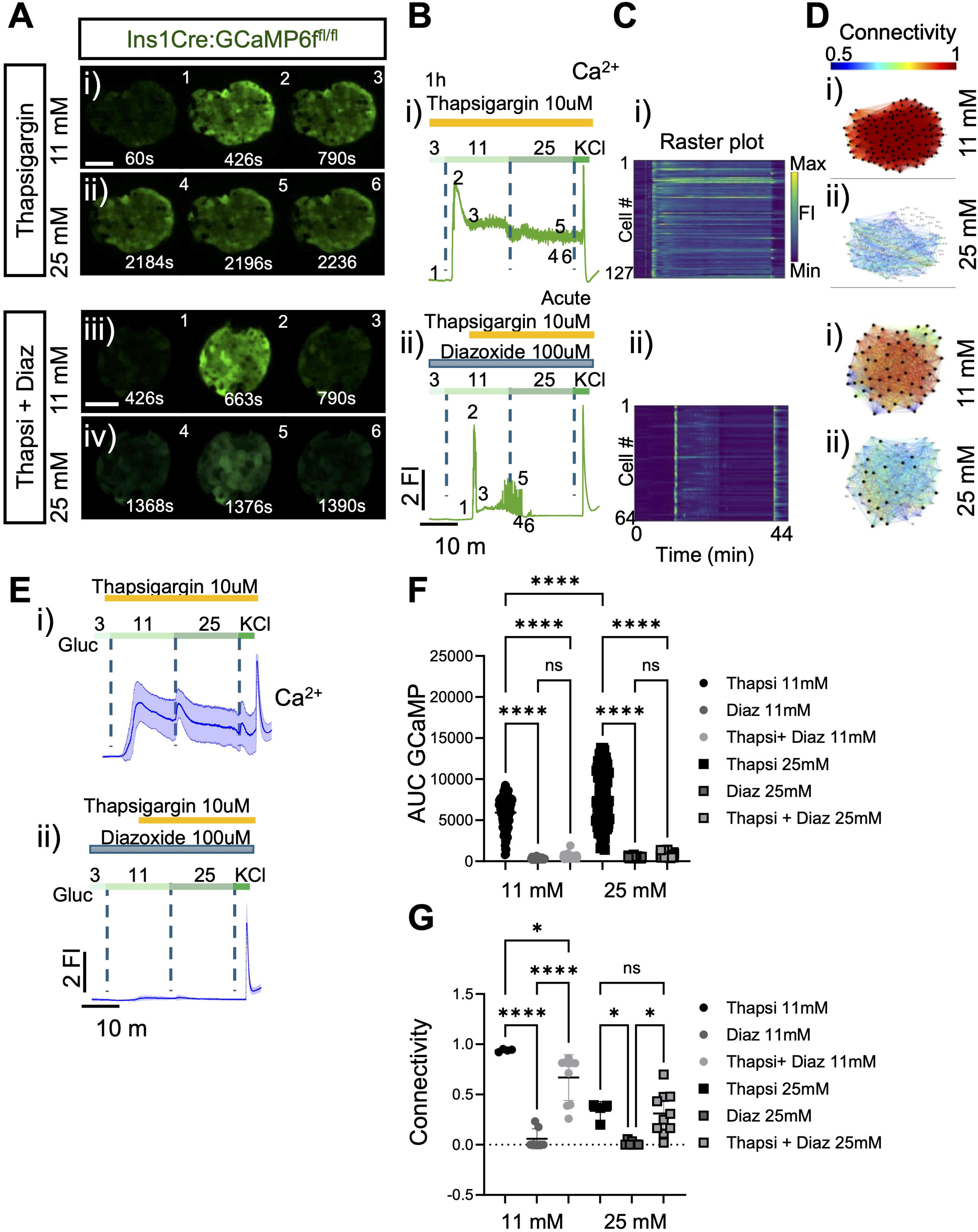
ER store depletion after SERCA pump inhibition with thapsigargin leads to sustained Ca^2+^ increases and lower connectivity. **A)** Series of confocal calcium imaging of pancreatic islet at the indicated time points (in sec.), exposed to 1hr preincubation with the SERCA pump inhibitor, thapsigargin and to diazoxide with acute exposure to thapsigargin during the imaging at **i)** 11 mM and **ii)** 25 mM glucose. Scale bar = 25 µm. **B)** Fluorescent traces of the GCaMP6 signal from the islets shown in **(A)** during the glucose ramp (3 mM, 11 mM, 25 mM) and membrane depolarization by 40 mM KCl. During **i)** 1h preincubation and **ii)** delivered after glucose stimulation. Of note, the acute delivery caused a transient calcium waves that eventually subsided. **C)** Raster plot corresponds to the signal from individual cells from the islets shown in **(A)**. **D)** Connectivity maps of the islets shown in **(A)** at **i)** 11 mM and **ii)** 25 mM glucose. **E)** Average calcium fluorescent traces from islets exposed to 1h thapsigargin or acutely exposed during a glucose ramp (3 mM, 11 mM, 25 mM) and membrane depolarization by 40 mM KCl. **F)** Calcium AUC quantifications from thapsigargin, diazoxide (Fig 3B-F) or acute thapsigargin+diazoxide at 11mM and 25mM glucose. Each symbol represents an islet (2-way paired ANOVA, Tukey’s correction \*\*\*\**P* ≤ .0001, ns, not significant; n=3 WT 115-islets for 1h thapsigargin, n=3 WT 45-islets for diazoxide and n=3 WT 58-islets from acute thapsigargin+diazoxide). **G)** Connectivity quantifications from 1h thapsigargin, diazoxide and acute thapsigargin+diazoxide at 11mM and 25mM glucose. Of note, during acute exposure to thapsigargin islets showed calcium activity which subsided over time. Each symbol represents an islet (2-way paired ANOVA, Tukey’s correction \**P* ≤ .05, \*\*\*\**P* ≤ .0001, ns, not significant; n=3 WT 4-islets for 1h thapsigargin, n=3 WT 7-islets for diazoxide and n=4 WT 10-islets from diazoxide+carbachol).

**Figure 5.** Interference with ER calcium store mobilization impairs the capability of the islets to generate sustained calcium waves. **A)** The normal function of ER calcium stores is required to transit from the first calcium transient to sustained calcium waves. **B)** Inhibition or hyperactivation of calcium mobilization from ER calcium stores interferes with calcium waves. Inhibition of ER calcium stores mobilization by the IP_3_R blocker 2-APB or depletion by the SERCA2 inhibitor thapsigargin interferes with calcium waves by decreasing overall calcium activity and decreasing connectivity. Conversely, hyperactivation of ER calcium stores mobilization by the muscarine receptor agonist carbachol, leads to an overall increase in single cell transients, however, the cells present uncoordinated calcium transients, which leads to low connectivity. Under normal conditions, ER calcium stores mobilization might allow beta-to-beta cell communication and increase their capability to synchronize their activity.

## 4. Discussion

The chief goal of the present work was to determine the extent to which intracellular Ca^2+^ release plays a role in the generation and propagation of Ca^2+^ waves across the islet, and the emergence of highly connected “hub” cells. We have assessed this in isolated primary islets in which Ca^2+^ changes are interrogated selectively in the beta cell using a molecularly targeted, genetic Ca^2+^ probe located exclusively in this cellular compartment. An important driver for the present work was a recent report from Postic and colleagues [42] showing that, in pancreatic slices, ryanodine receptor activation or inactivation, respectively, blocked and slowed, or promoted, Ca^2+^ release events under different conditions. The sources of released calcium accessed by ryanodine receptors is, however, unclear, and may include (or even be restricted to) acidic intracellular compartments including secretory granules [43]. This question was not explored by Postic et al [42]. In deploying a non-molecularly-targeted calcium probe, the sub-cellular and cellular provenance of the recorded signals cannot be assigned unambiguously in the latter report, and contamination with events in other endocrine cell types is not ruled out.

We have therefore combined here the use of a targeted calcium sensor, GCaMP6f, expressed exclusively in beta cells under the control of an Ins1*Cre* knock-in allele [19] alongside three complementary pharmacological approaches to explore the importance of ER calcium stores in glucose-induced Ca^2+^ waves across the islet. We assessed firstly the role of IP_3_R, either through the inhibition of these receptors with 2-APB or their activation after IP_3_ generation in response to muscarinic stimulation. We acknowledge that the former, non-selective drug exerts pleiotropic effects which go beyond IP_3_R inhibition. These include depletion of the ER Ca^2+^ store through the combined action as an inhibitor of SERCA pumps and by activating an ER Ca^2+^ leak channel (see Results, 3.1) [39–41]. Despite this limitation, this tool allowed us to explore the role of intracellular Ca^2+^ mobilization on glucose-stimulated oscillations and waves in the islet. Likewise, we note that carbachol treatment elicits effects beyond IP_3_ generation, including the production of diacylglycerol and the activation of protein kinase C family members. However, these events are likely to have minimal impact on intracellular Ca^2+^ dynamics over the time course of the present experiments (minutes), though actions on intercellular connectivity cannot be ruled out.

We have deployed a protocol involving an initial challenge with an intermediate (11 mM) glucose concentration which was then followed by an increase in glucose to 25 mM. Our analyses of Ca^2+^ dynamics at each glucose concentration thus allowed us to explore the effects of pharmacological manipulations on both the responses to an initial large rise in glucose (3-11 mM) and then to a second increase (11-25mM). It is therefore important to note that the responses in each case were distinct, with an initial, highly coordinated response to the first challenge (11 mM) which in control conditions then transitioned to baseline oscillations which often traversed the islet (Fig. 1 b,c). In contrast, the response to the further increase in glucose to 25 mM usually involved a dip in average Ca^2+^ levels which was then followed by Ca^2+^ oscillations above an elevated baseline. This difference in the temporal properties of the responses must therefore be borne in mind when considering the impacts of pharmacological treatments at each glucose concentration.

Interestingly, 2-APB had clear effects on overall Ca^2+^ increases in response to 11 mM glucose, with integrated calcium increase and oscillation peak amplitude showing highly significant decreases, or a tendency towards a decrease, respectively. In contrast, at the higher glucose concentration of 25 mM, both the overall calcium response and the properties of individual Ca^2+^ peaks were substantially diminished by the drug (Fig. 2). These observations suggest that intracellular Ca^2+^ stores may become more important when Ca^2+^ influx through VGCCs is maximally stimulated. Accordingly, whilst cell-cell connectivity was similar at 25 and 11 mM glucose in the absence of further additions, 2-APB caused a substantially weakening of connectivity at the higher concentration of glucose, further supporting a role for intracellular Ca^2+^ in supporting beta cell-beta synchrony under hyperglycemic conditions. Taken together, these findings support the view that ER Ca^2+^ plays a non-negligible role in modulating glucose-induced cytosolic Ca^2+^ changes in response to glucose.

We show next that intracellular IP_3_ generation in response to carbachol potentiates glucose-induced Ca^2+^ increases and has little effect om cell-cell connectivity at 11 mM glucose. By using diazoxide to hyperpolarise the plasma membrane and suppress Ca^2+^ influx through VGCCs, we also show that the effects of this muscarinic agonist to prompt intracellular Ca^2+^ oscillations and waves are strongly dependent on Ca^2+^ influx through VGCCs, with the calcium oscillations under these conditions being poorly connected between cells. Thus, Ca^2+^ oscillations driven purely, or very largely, by intracellular Ca^2+^ release channels have little capacity to instigate and support Ca^2+^ oscillations which can traverse the islet.

Nevertheless, we did note that under these conditions a small number of cells displayed higher connectivity, resonant of “small worlds” behaviour. As noted recently [20] the recording of Ca^2+^ dynamics with GCaMP6, as opposed to the earlier use of non-targeted small molecule chemical probes (e.g. fluo-4) [18], leads us to conclude that the majority of beta cells enjoy a high degree of connectivity with other beta cells (Figure 4 B). Thus, “small worlds” behaviour, involving a small subset of unusually highly-connected beta cells is relatively rare in the healthy islet. Nonetheless, a gradient of “connectedness” exists, with subpopulations of more highly connected cells identifiable. Future studies will be required to determine to what extent these populations were affected by manipulation of ER calcium stores.

As a final approach, we show that cell-cell coordination is still possible after the complete depletion of ER stores with the SERCA pump inhibitor, thapsigargin [44]. However, and in common with the approaches above, the effects of this drug to impede Ca^2+^ dynamics and connectivity were more marked at 25 mM than 11 mM glucose.

## Conclusions

The present findings provide evidence for a limited, but non-negligible and glucose concentration-dependent role for ER calcium stores in the maintenance and transmission of glucose-induced Ca^2+^ waves across the beta cell syncytium of the islet (Graphical Abstract). Thus, we show that ER store filling and mobilisation need to be poised in order to ensure optimal Ca^2+^ dynamics and wave transmission, likely to be associated with appropriately regulated insulin secretion. The requirement for internal stores assumes greater importance at high glucose levels which are likely to pertain in (pre)diabetes. Control of beta cell Ca^2+^ mobilisation may thus be of potential therapeutic value to maintain beta cell-beta connectivity and insulin secretion in these circumstances, and may even slow or prevent disease progression

## Acknowledgements

*Funding.* G.A.R. was supported by Wellcome Trust Senior Investigator (WT098424AIA) and Investigator (WT212625/Z/18/Z) Awards, and MRC Programme grant (MR/R022259/1), Diabetes UK (BDA 16/0005485), an NIH-NIDDK project grant (R01DK135268) a CIHR-JDRF Team grant (CIHR-IRSC TDP-186358 and JDRF 4-SRA-2023-1182-S-N), CRCHUM start-up funds, and an Innovation Canada John R. Evans Leader Award (CFI 42649). LD was support by a CIHR/IRSC Post-doctoral Fellowship (#489982).

*Duality of Interest.* G.A.R. has received grant funding from Sun Pharmaceuticals Inc. and Laboratoires Servier and is a consultant for Sun Pharmaceuticals Inc. No other potential conflicts of interest relevant to this article were reported.

## 5. CRediT author statement

**LD.** conceptualisation, methodology, analysis: **KD**, investigation, analysis, writing; **GR**, conceptualisation, writing, supervision

