## Supplementary figures for "ER calcium stores contribute to glucose-induced Ca^2+^ waves and intercellular connectivity in mouse pancreatic islets"

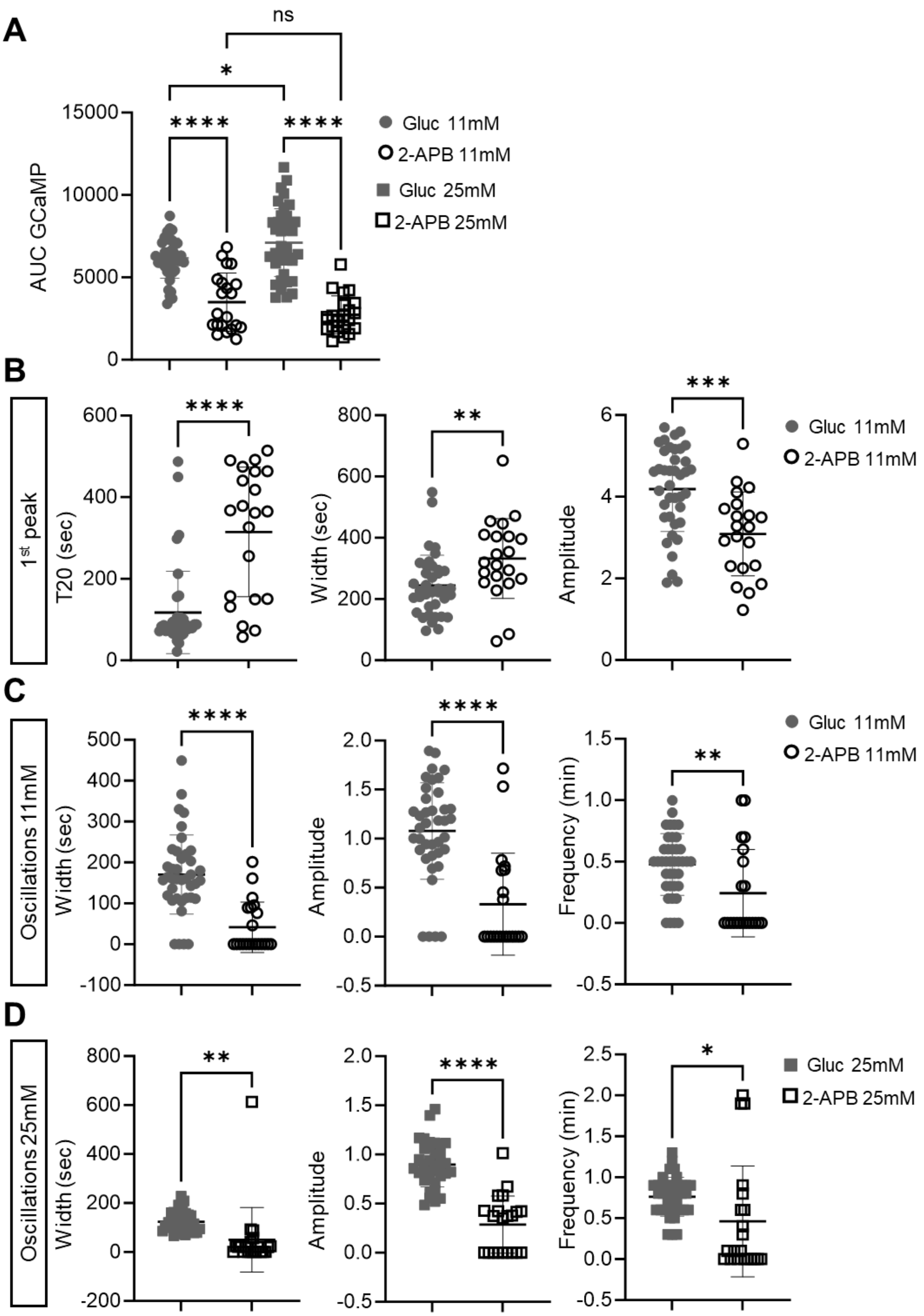

**Supplementary Figure 1. $IP_3R$ inhibition by 2-APB impairs $Ca^{2+}$ dynamics and connectivity. A)** Calcium AUC quantifications from glucose and 2-APB (Fig 2) islets at 11mM and 25mM glucose. **B)** $T_{20}$ is the time taken from the glucose addition in the media to present an increase of 20% over the baseline. The amplitude of the 1st peak is defined as the maximum value recorder during the 1st increase. The peak width is defined as the time spent with an intensity >20% over the base line. The graphs comparing the $T_{20}$, peak width and peak amplitude between glucose and 2-APB exposed islets. **C)** The frequency analysis at 11mM and 25mM covering the last 10 min of each section. Average peak width, peak amplitude and peak frequency quantifications of islets incubated with glucose and 2-APB at 11mM. **D)** Average peak width, peak amplitude and peak frequency quantifications of islets incubated with glucose and 2-APB at 25mM. Each symbol represents an islet (2-way paired ANOVA, Tukey's correction **P* ≤ .05, ***P* ≤ .01, *****P* ≤ .0001, ns, not significant; n=3 WT 39-islets for glucose and n=3 WT 21-islets for 2-APB. Data are mean ± SD.)

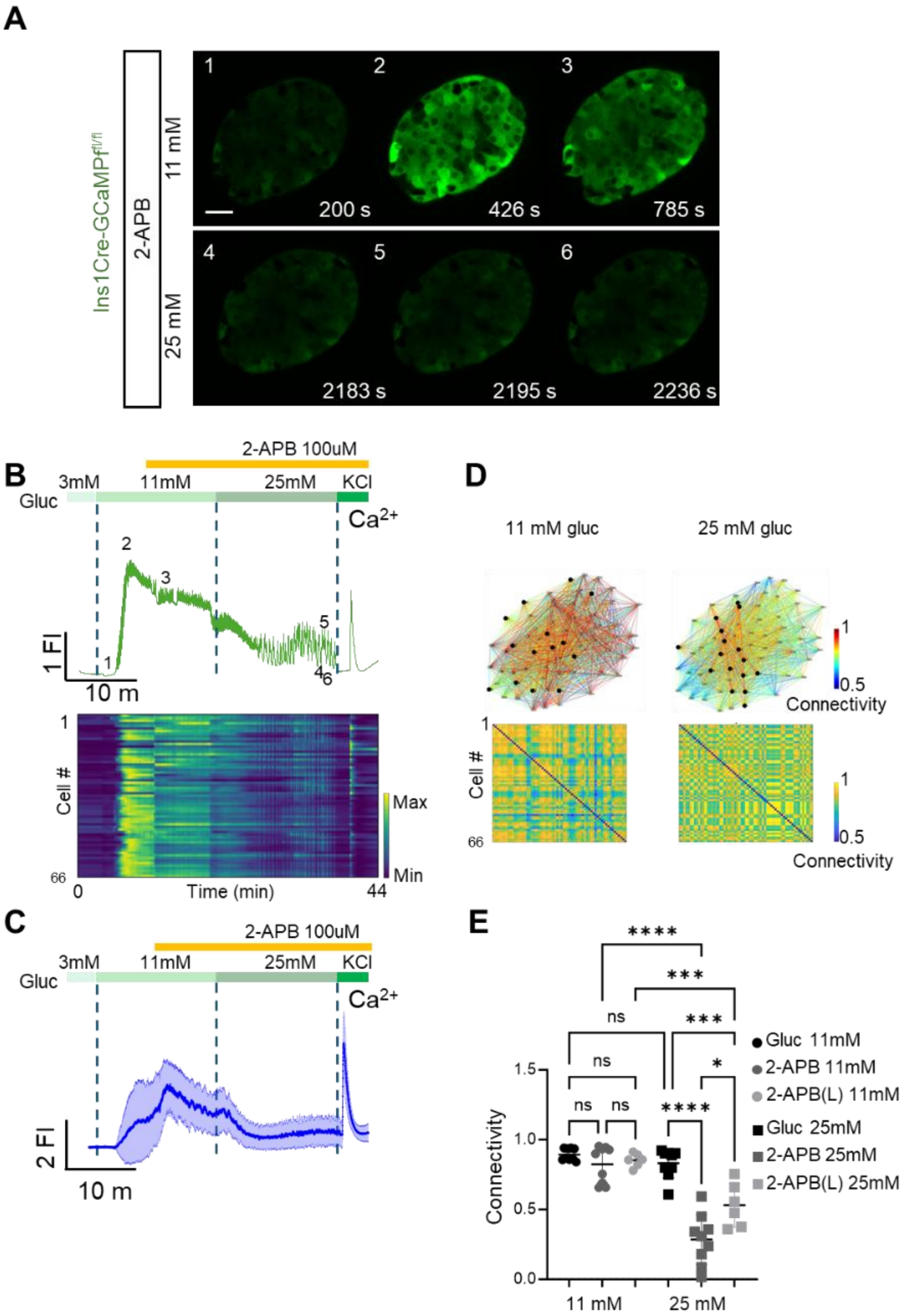

A
Ins1Cre-GCaMP$^{fl/fl}$
2-APB
11 mM
25 mM
1
2
3
200 s
426 s
785 s
4
5
6
2183 s
2195 s
2236 s
B
2-APB 100uM
3mM
11mM
25mM
KCl
Gluc
Ca$^{2+}$
1 FI
10 m
Cell #
Max
Min
Time (min)
0
44
66
C
2-APB 100uM
3mM
11mM
25mM
KCl
Gluc
Ca$^{2+}$
2 FI
10 m
D
11 mM gluc
25 mM gluc
Connectivity
Cell #
66
E
Connectivity
11 mM
25 mM
Gluc 11mM
2-APB 11mM
2-APB(L) 11mM
Gluc 25mM
2-APB 25mM
2-APB(L) 25mM
ns
****
***
*

**Supplementary Figure 2. Inhibition of $IP_3R$ by 2-APB impairs glucose-induced intracellular $Ca^{2+}$ dynamics and connectivity. A)** Series of confocal calcium imaging of pancreatic islet at the indicated time points (in sec.), exposed to the IP3-inhibitor 2-Aminoethoxydiphenyl borate (2-APB) at **i)** 11 mM and **ii)** 25 mM glucose. Scale bar = 25 µm. The 2-APB was delivered after 10min of glucose increase from 3mM to 11mM. **B)** Fluorescent trace of the GCaMP6 signal from the islet shown in **(A)** during the glucose ramp (3 mM, 11 mM, 25 mM) and membrane depolarization by 40 mM KCl. The raster plot corresponds to the signal from individual cells. **C)** Average calcium fluorescent traces from islets exposed to 2-APB during a glucose ramp (3 mM, 11 mM, 25 mM) and membrane depolarization by 40 mM KCl. The blue line shows the average GCaMP6f fluorescent and the light-blue the standard deviations from the imaged islets (n=4 WT 103-islets). **D)** Connectivity map and matrix of the islet shown in **(A)** at **i)** 11 mM and **ii)** 25 mM glucose. **E)** Connectivity from individual islets glucose, 2-APB from the beginning of the recording or later addition 2-APB(L) (10m after glucose increase from 3mM to 11mM glucose). (2-way paired ANOVA, Tukey’s correction **P* ≤ .05, ****P* ≤ .001, *****P* ≤ .0001, ns, not significant; glucose n=4 WT 9-islets; 2-APB n=4 WT 9-islets; 2-APB (Late) n=3 WT 6-islets).

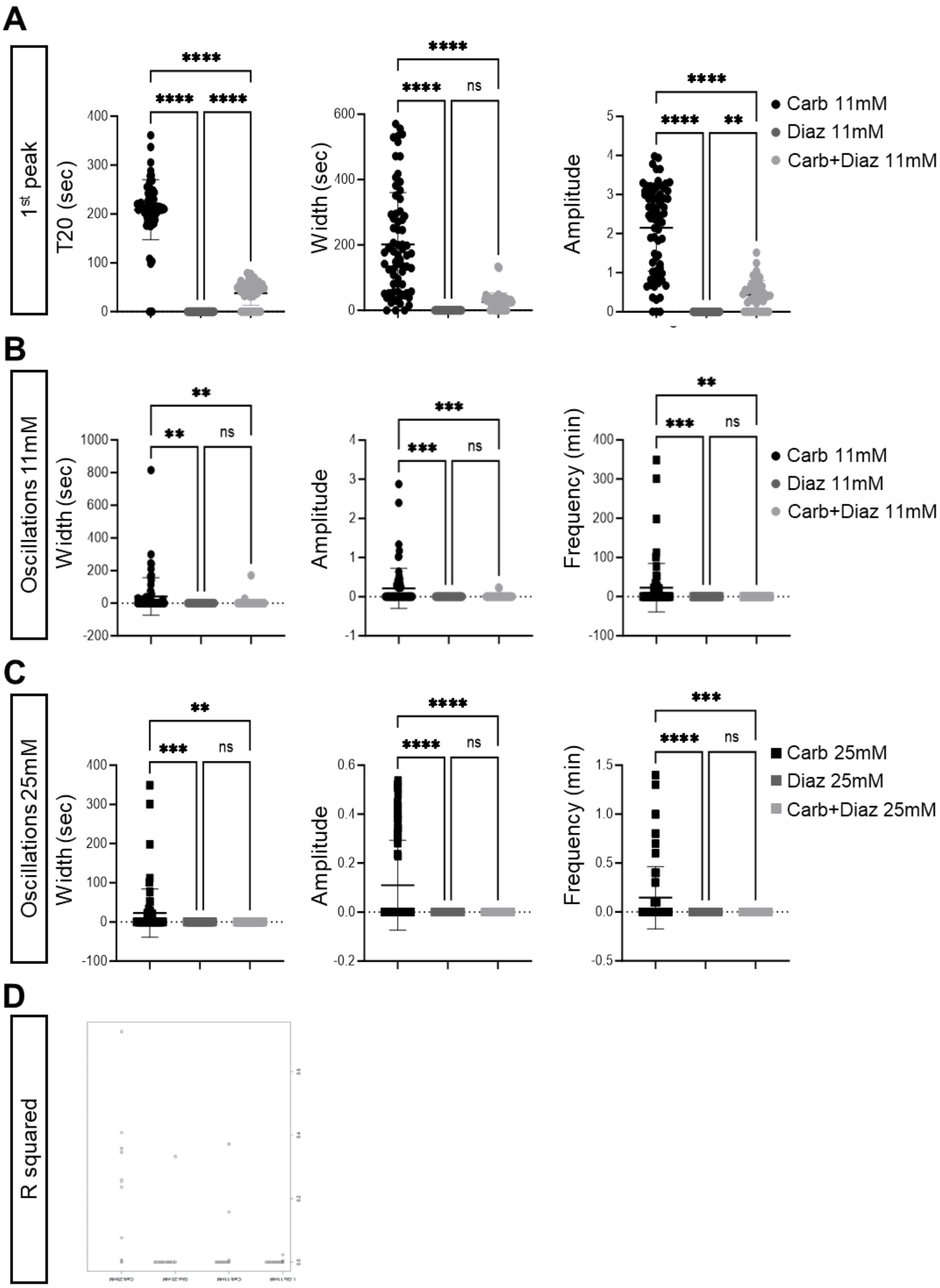
A
1st peak
T20 (sec)
Width (sec)
Amplitude
Carb 11mM
Diaz 11mM
Carb+Diaz 11mM
B
Oscillations 11mM
Width (sec)
Amplitude
Frequency (min)
Carb 11mM
Diaz 11mM
Carb+Diaz 11mM
C
Oscillations 25mM
Width (sec)
Amplitude
Frequency (min)
Carb 25mM
Diaz 25mM
Carb+Diaz 25mM
D
R squared

**Supplementary Figure 3. Inhibition of $IP_3R$ by Carbachol impairs $Ca^{2+}$ dynamics and connectivity. A)** The graphs comparing the $T_{20}$, peak width and peak amplitude between carbachol, diazoxide and carbachol+diazoxide exposed islets. **B)** The frequency analysis at 11mM and 25mM covering the last 10 min of each section. Average peak width, peak amplitude and peak frequency quantifications of islets incubated with carbachol, diazoxide and carbachol+diazoxide at 11mM. **C)** Average peak width, peak amplitude and peak frequency quantifications of islets incubated with carbachol, diazoxide and carbachol+diazoxide at 25mM. Each symbol represents an islet (2-way paired ANOVA, Tukey's correction *$P \leq .05$, **$P \leq .01$, ***$P \leq .001$, ****$P \leq .0001$, ns, not significant; n=3 WT 71-islets for carbachol, n=3 WT 45-islets for diazoxide and n=3 WT 47-islets from diazoxide+carbachol. Data are mean ± SD.)

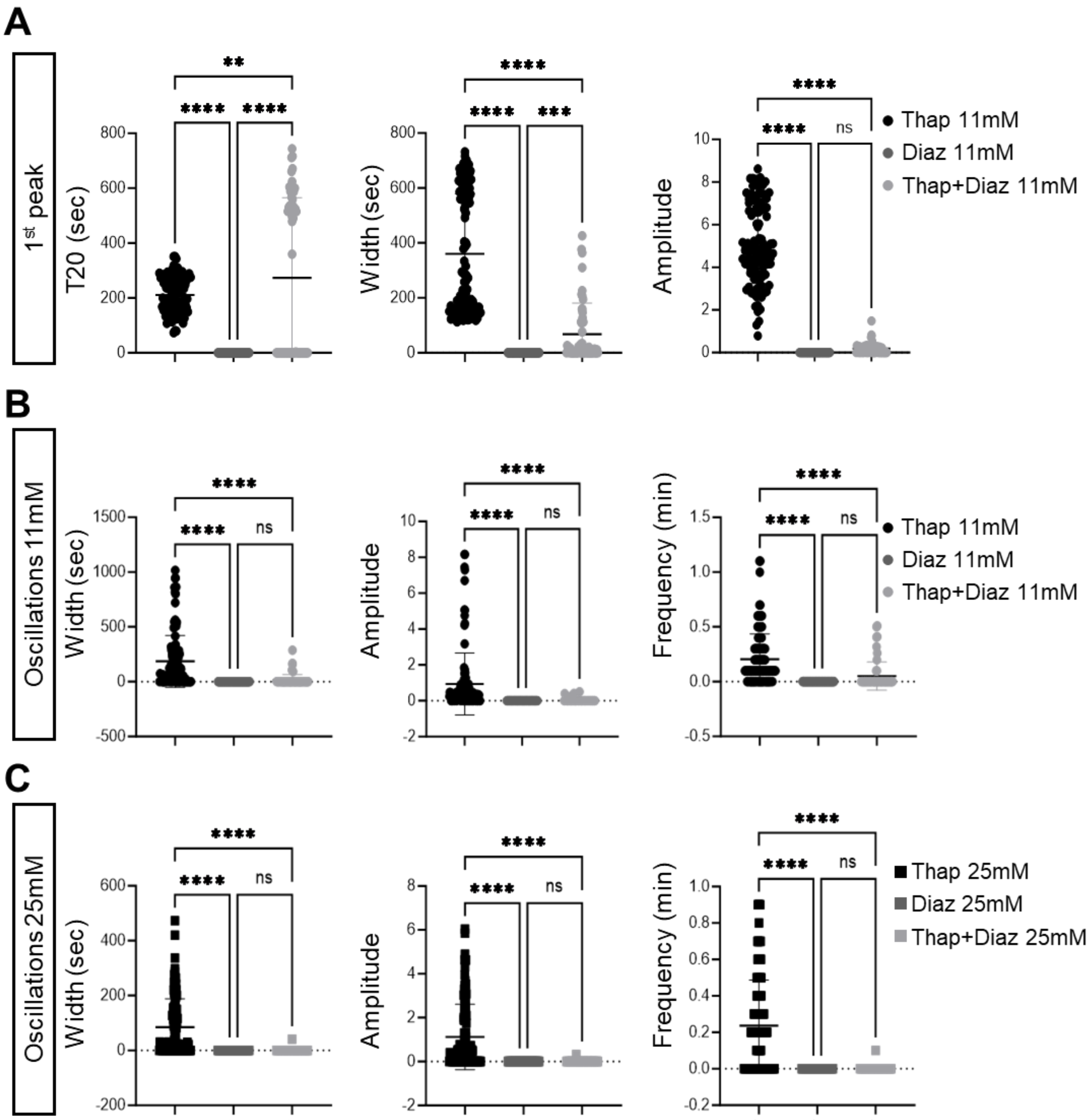
A
1st peak
T20 (sec)
Width (sec)
Amplitude
**
****
***
ns
Thap 11mM
Diaz 11mM
Thap+Diaz 11mM
B
Oscillations 11mM
Width (sec)
Amplitude
Frequency (min)
C
Oscillations 25mM
Width (sec)
Amplitude
Frequency (min)
Thap 25mM
Diaz 25mM
Thap+Diaz 25mM

**Supplementary Figure 4. Inhibition of SERCA2 by thapsigargin impairs $Ca^{2+}$ dynamics and connectivity. A)** The graphs show the $T_{20}$, peak width and peak amplitude between 1h thapsigargin, diazoxide and acute thapsigargin+diazoxide exposed islets. **B)** The frequency analysis at 11mM and 25mM covering the last 10 min of each section. Average peak width, peak amplitude and peak frequency quantifications of islets incubated with 1h thapsigargin, diazoxide and acute thapsigargin+diazoxide at 11mM. **D)** Average peak width, peak amplitude and peak frequency quantifications of islets incubated with 1h thapsigargin, diazoxide and acute thapsigargin+diazoxide at 25mM. Each symbol represents an islet (2-way paired ANOVA, Tukey's correction $^{*}P \leq .05$, $^{**}P \leq .01$, $^{***}P \leq .001$, $^{****}P \leq .0001$, ns, not significant; n=3 WT 71-islets for carbachol, n=3 WT 115-islets for 1h thapsigargin, n=3 WT 45-islets for diazoxide and n=3 WT 58-islets from acute thapsigargin+diazoxide. Data are mean ± SD.)
